# A general method to quantify ligand-driven oligomerization using single- or two-photon excitation microscopy

**DOI:** 10.1101/477307

**Authors:** Michael R. Stoneman, Gabriel Biener, Richard J. Ward, John D. Pediani, Dammar Badu, Ionel V. Popa, Graeme Milligan, Valerică Raicu

## Abstract

Current technologies for probing membrane protein association and stability in cells are either very laborious or lack the bandwidth needed for fully quantitative analysis. Here we introduce a platform, termed *one-* or *two-dimensional fluorescence intensity fluctuation spectrometry*, for determining the identity, abundance, and stability of oligomers of differing sizes. The sensitivity of this approach is demonstrated by using monomers and oligomers of known sizes in both solutions and cell membranes. The analysis was extended to uncover the oligomeric states and their stability for both the epidermal growth factor receptor, a receptor tyrosine kinase, and the G protein-coupled secretin receptor. In both cases, agonist ligand binding shifted the equilibrium from monomers or dimers to rather large oligomers. Our method can be used in conjunction with a variety of light-based microscopy techniques, is several orders of magnitude faster than current approaches, and is scalable for high-throughput analyses.

---

The association of membrane proteins into dimers or higher-order oligomers is thought to regulate biological function. For example, the lateral association of receptor tyrosine kinases (RTKs) into oligomers controls RTK activation, as the proximity of two kinase domains in the oligomers is required for their cross-phosphorylation^1–3^. Likewise, the association of G protein-coupled receptors (GPCRs)^4–6^ into oligomers is believed to affect their interactions with G-proteins and other effector molecules^7–11^. Defects in the association of cell surface receptors have been linked to human diseases^12–14^. Yet, the discrimination of the oligomeric state of a membrane receptor in live cells remains a significant experimental challenge, with different studies from different laboratories often producing contradicting results^15,16^. Whether such contradictions stem from data over-interpretation or built-in structural and functional versatility of these receptors is yet to be clarified^17,18^.

Existing technologies either are laborious and therefore slow or lack the capability of discriminating between different oligomeric sizes. For example, Förster Resonance Energy Transfer (FRET) is potent at extracting the geometry and stoichiometry of protein complexes^3,19,20^ but relies on sophisticated analyses that are not easily scalable to probing receptor oligomerization under different experimental conditions or to fast screening of ligands for their therapeutic potential. In addition, FRET requires two labels and faces difficulties for inter-protomer distances larger than a spectroscopic quantity called Förster’s radius. Fluorescence fluctuation spectroscopies^21–25^, such as Number and Brightness analysis (N&B) and Spatial Intensity Distribution Analysis (SpIDA), are comparatively simpler but currently only provide average values of the oligomer size and concentration over entire populations of oligomers with different concentrations and sizes. Because of this, a mixture of, e.g., 33 % tetramers and 67 % monomers would appear on average as dimers. In addition, inhomogeneously distributed concentrations may inadvertently appear as oligomer size distributions. Clearly, methods for probing association of cellular receptors in cells with accuracy and speed are needed for a better understanding of receptors function and for the development of effective therapies that target protein-protein interactions.

We have developed a method for determination of the identity, abundance, and stability of differing quaternary structures formed by transmembrane receptors in living cells under various environmental conditions. Our method, termed *one-* or *two-dimensional fluorescence intensity fluctuation spectrometry* (i.e., FIF or 2D-FIF spectrometry, respectively), like N&B and SpIDA, exploits the fact that fluctuations in the measured fluorescence intensity are proportional to the number of fluorescent molecules associated with one another and therefore passing together through a light beam used for their excitation. Unlike other methods, however, 2D FIF extracts critical information also from inherent variations in the molecular brightness and concentration distributions within the regions of interest. These same variations constitute the main limitation and source of difficulties for other fluorescence intensity fluctuation-based methods. In addition, 2D FIF may be implemented on various fluorescence-based microscopes, including confocal, two-photon excitation, and total internal reflection microscopes, and will be therefore accessible to most research or pharmaceutical laboratories. We have implemented the 2D FIF analysis into a computer program (submitted with the manuscript) that produces a complete set of data per sample within less than one day using a standard PC. Currently, 2D FIF is orders of magnitude faster than FRET spectrometry^26^ (which, nevertheless, also provides relative distances between protomers), and it may be accelerated further by using automated image acquisition systems and faster computers; it is therefore scalable to high-throughput screening of natural and artificial ligands that can shift the monomer-dimer-oligomer equilibrium.

Our method starts from imaging cells expressing fluorescently-labeled proteins of interest, which can be done, for example, with a standard confocal microscope. Large regions of interest (ROI) are then demarcated using an ROI-selection tool, and a dedicated computer program generates small ROI segments using a mathematical procedure described in the Methods section. The ROI segmentation procedure not only makes it possible to convert local intensity variations within the ROI into useful information, but it also speeds up the analysis process, by generating over ten thousand ROI segments (i.e., brightness and concentration data points) from just hundreds or fewer ROIs.

The ROI selection and segmentation process is illustrated in Figure 1 (panels a and b) using confocal images of the basolateral membrane of cells expressing a monomeric enhanced green fluorescent protein (mEGFP) construct associated to the plasma membrane via a lipidated peptide anchor (PM-1-mEGFP, see Methods). A fluorescence intensity distribution is generated for each ROI segment (Figure 1b inset), from which the average intensity as well as the variance of the distribution are computed. A one-dimensional brightness spectrogram is then computed for each ROI segment, as shown in Figure 1c. Analysis of this spectrogram gives the effective molecular brightness of a monomer (or protomer, 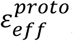). A similar analysis was performed for an equivalent tandem mEGFP construct, PM-2-mEGFP, in which two molecules of mEGFP are linked together (Figure 1d). Here the effective brightness peak was located roughly at double the effective brightness of the monomeric construct. Reassuringly, these results were consistent with those obtained when using purified monomeric and tandem constructs of YFP in solution as well as from the monomeric and dimerizing EGFP constructs in cell membranes using a two-photon microscope (see Supplementary Figure 1). The existence of a small dimeric component in the PM-1-mEGFP spectrogram and oligomeric components in the PM-2-mEGFP spectrogram is probably due to weak non-specific interactions.

**Figure 1.**
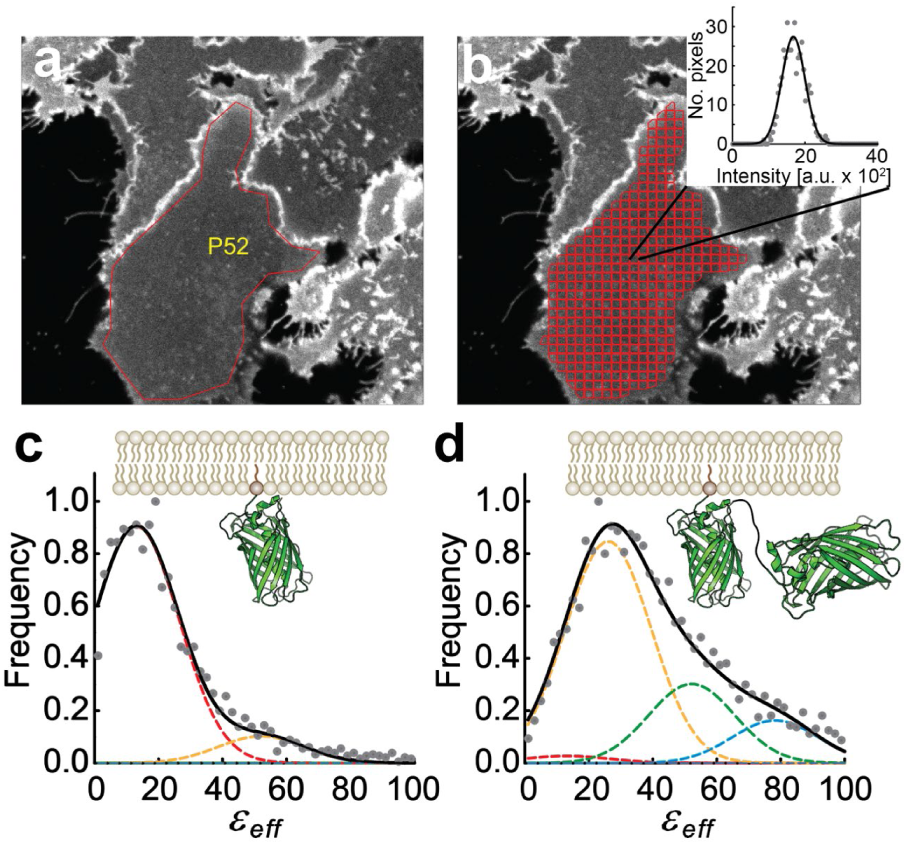
Illustration of the data reduction process in two-dimensional Fluorescence Fluctuation (2D-FIF) Spectrometry using single-photon excitation. **a,** Typical fluorescence image of Flp-In™ T-REx™ 293 cells expressing a plasma membrane-targeted mEGFP construct (PM-1-mEGFP). The overlaid polygon (P52) indicates a region of interest (ROI) comprising a patch of the basolateral membrane of a cell. **b**, Software-generated image segmentation of the ROI in **a** using the moving squares method (see Methods). Inset, fluorescence intensity histogram (circles) of a single image segment, together with its Gaussian curve fit (solid line). The mean and width of the Gaussian are used to calculate the brightness (*ε*_*eff*_) and concentration for each segment (see Methods). **c-d**, Normalized frequency distribution assembled from **(c)** 3,582 and **(d)** 4,185 total *ε*_*eff*_ values obtained from monomeric (PM-1-mEGFP) or tandem (PM-2-mEGFP) mEGFP constructs was fit to a sum (solid black curves) of Gaussian functions (represented by dashed lines with various colors, to find the brightness of single mEGFP protomers, 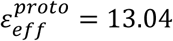. Each Gaussian peak position was set to a value of 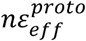, where *n* is the number of protomers in an oligomer. The *ε*_*eff*_ distribution for PM-1-mEGFP is primarily comprised of monomers (dashed, red Gaussian curve), with its peak value positioned at 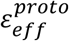, while the PM-2-mEGFP spectrogram is mostly captured by a dimer model (dashed, yellow Gaussian curve), with its peak positioned at 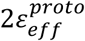.

We used the epidermal growth factor receptor (EGFR) fused to mEGFP as a testbed for our method, as this receptor’s oligomerization status and functional role in the presence and absence of ligands are well-understood^1,2,27^. Wild-type EGFR presents ligand-dependent oligomerization, while a mutant form containing a pair of domain II mutations (Tyr^251^Ala and Arg^285^Ser) is unable to oligomerize^28^, either in the presence or absence of ligand. Following the same process described in Figure 1 for confocal imaging and data analysis of monomeric and tandem (dimeric) fluorescent constructs, we obtained the brightness (*ε*_*eff*_) and average intensity for each ROI segment of cell membranes containing mEGFP-EGFR. Dividing the average intensity by the brightness of the monomeric construct (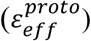) derived above and the volume of the laser beam focal volume (see Methods), we obtained the concentration of EGFR protomers in each ROI segment.

The frequency of occurrence of each effective brightness-protomer concentration pair was represented as two-dimensional surface plots in Figure 2 for the wild-type EGFR, both in the presence and absence of its canonical ligand, the epidermal growth factor (EGF). For wild-type EGFR, the distribution of oligomer sizes was centered around dimers for untreated cells (Figure 2a_2_,a_3_) and tetramers for ligand-treated cells (Figure 2b_2_,b_3_) and dramatically shifted towards larger oligomer sizes as the concentration of the receptor increased. By contrast, for the oligomerization-impaired mutant EGFR, the oligomer size remained unchanged after addition of EGF as ligand (Supplementary Figure 3). Rather gratifyingly, all of these findings are in agreement with recent literature^1,2,27^.

**Figure 2.**
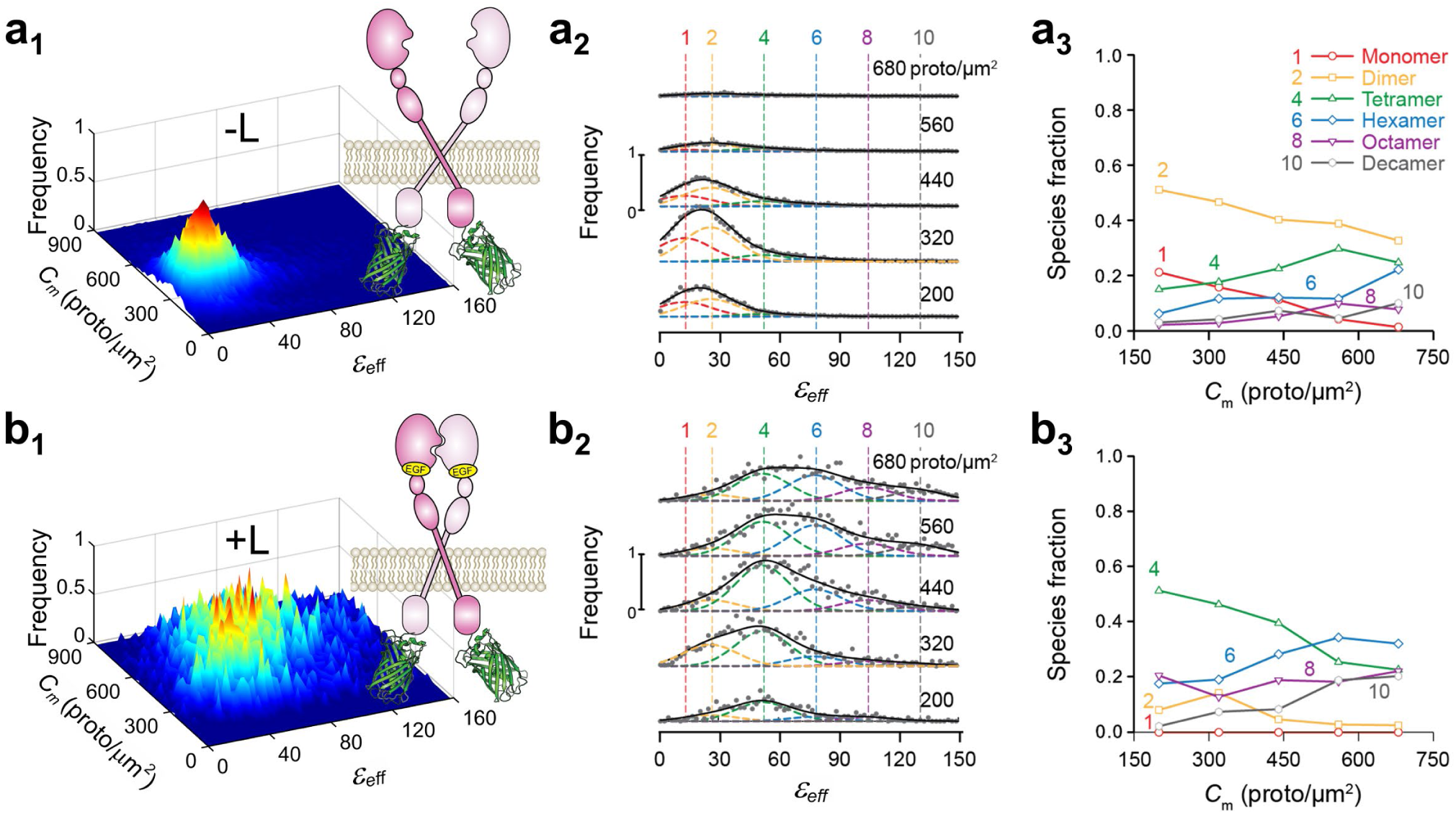
Analysis of effective brightness (*ε*_*eff*_) distributions obtained from single-photon excitation of cells expressing wild-type EGFR in the absence of ligand (-L) or after ten-minute treatment with 100 nM ligand (+L). a1-b1 (column 1), Frequency of occurrence of *ε*_*eff*_ for each protomer concentration using **(a1)** 25,740 and **(b1)** 6,812 total ROI segments to construct the distribution. **a2-b2 (column 2),** Cross sections through the surface plots in panels **a1** and **b1** for different total concentration ranges; average concentration for each range (in protomer/μm^2^) is indicated above each plot. The vertical dashed colored lines indicate the peak positions for the brightness spectra of monomers, dimers, etc., obtained from (or predicted based on) the simultaneous fitting of the PM1- and PM2-mEGFP spectrograms, which were used as standards of brightness in the analysis (see below). **a3-b3 (column 3),** Relative concentration of protomers in each oligomeric species vs. total protomer concentration, as derived by decomposing the spectrograms in column 2 into Gaussian components. The *ε*_*eff*_ distribution for each concentration range was fitted with a sum of six Gaussians; the peak of each Gaussian was set to 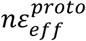, where *n* is the number of protomers in a given oligomer (e.g., 1, 2, 4, etc.), while the standard deviations (SD) were fixed at 13.4 (a.u.). The peak positions and their SDs were obtained from measurements on cells expressing PM-1-mEGFP or PM-2-mEGFP (see Figure 1). Only the Gaussian amplitudes (*A*_*n*_) were adjusted in the process of data fitting in **b**, which gave the fraction of protomers (shown in **column 3**) for each oligomeric species, i.e., *n*_*i*_*A*_*i*_/∑_*n*_ *nA*_*n*_.

We next used 2D FIF spectrometry to investigate the oligomerization of the secretin receptor (SecR)^29^, a class B GPCR whose oligomerization behavior, like that of other GPCRs, is not fully understood. While several experiments have indicated that GPCRs form functional homo- or hetero-oligomeric complexes *in vivo* as well as *in vitro*^18,24,30–34^, there have been suggestions that not all GPCRs are multimeric or that oligomerization is not essential for function^35–37^. Although some of the discrepancies are likely related to the variety of functions, and hence behaviors, of these receptors, some may be traced back to the inadequacies of some of the methods used.

We used again a confocal microscope to image cells expressing SecR-mEGFP fusion proteins in their plasma membrane, and found that the 2D FIF spectrogram was significantly broader for ligand-treated cells compared to untreated cells (compare panels a1 and b1 in Figure 3); this was true for all SecR expression levels investigated (Figure 3 panels a2 and b2). While the dominant species consisted of monomers in the case of untreated cells, the dimer fraction was significantly larger for secretin-treated cells, with the entire distribution being shifted towards larger sizes (compare panels a3 and b3 in Figure 3). A plausible explanation for this behavior is that SecR presents two different interfaces that allow it to bind laterally to other SecR molecules: one which allows monomers to dimerize and another one that permits association of dimers into higher-order oligomers^32^. The receptors shuttle between the associated and unassociated states, with the residence time in each state depending on concentration (see Figure 3a3). Ligand binding stabilizes the monomer-monomer interface, causing the dimer to dominate at any of the concentrations investigated (see Figure 3b3), thereby allowing higher order oligomers to form via the second binding interface.

**Figure 3.**
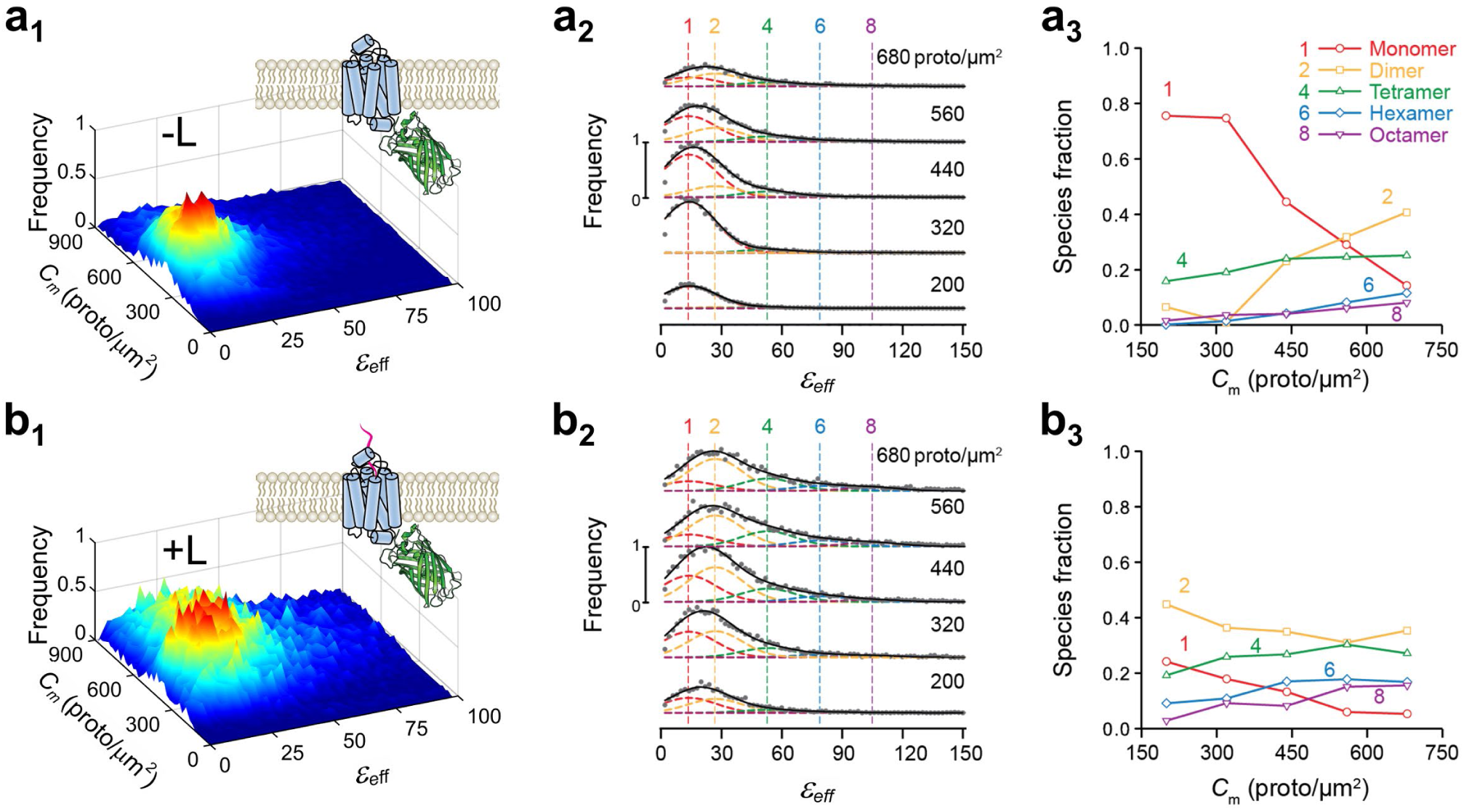
Analysis of effective brightness (*ε*_*eff*_) distributions obtained from single-photon excitation of cells expressing wild-type secretin receptor in the absence of ligand (-L) or after ten-minute treatment with 100 nM ligand (+L). a1-b1 (column 1), Frequency of occurrence of *ε*_*eff*_ for each protomer concentration using **(a1)** 64,619 and **(b1)** 29,839 total segments to construct the distribution. **a2-b2 (column 2),** Stacks of cross sections through the surface plots in panels **a1** and **b1**; average concentration for each range (in protomer/μm^2^) is indicated above each plot. Vertical dashed lines indicate peak positions for the brightness spectra of monomers, dimers, etc., obtained from (or predicted from) the simultaneous fitting of the PM-1- and PM-2-mEGFP spectrograms used as standards of brightness (see caption to Figure 2). **a3-b3 (column 3),** Relative concentration of protomers within each oligomeric species vs. total concentration of protomers, as derived from unmixing of the curves in column 2 into different Gaussian components. Samples were as follows: wild-type secretin receptor treated with vehicle (-L) (**row a** of graphs) or secretin (+L) for 10 minutes (**row b**). All analysis followed the method described in the caption to Figure 2.

We next fixed the cells after 10 and 30 minutes, respectively, of treatment with 100 nM secretin to test whether SecR oligomerization depends on the duration of the treatment. In addition, this time we used a two-photon microscope for imaging, to test whether the results are independent of the imaging modality. As seen in Figure 4, the results obtained for untreated and 10-minute treated cells were consistent with those obtained before with the confocal microscope, which proves the instrument-independent character of this method. In addition, prolonged treatment with ligand completely abolished the monomeric form of the receptor, with tetramers and even hexamers dominating over a broad range of concentrations (see Figure 4c3).

**Figure 4.**
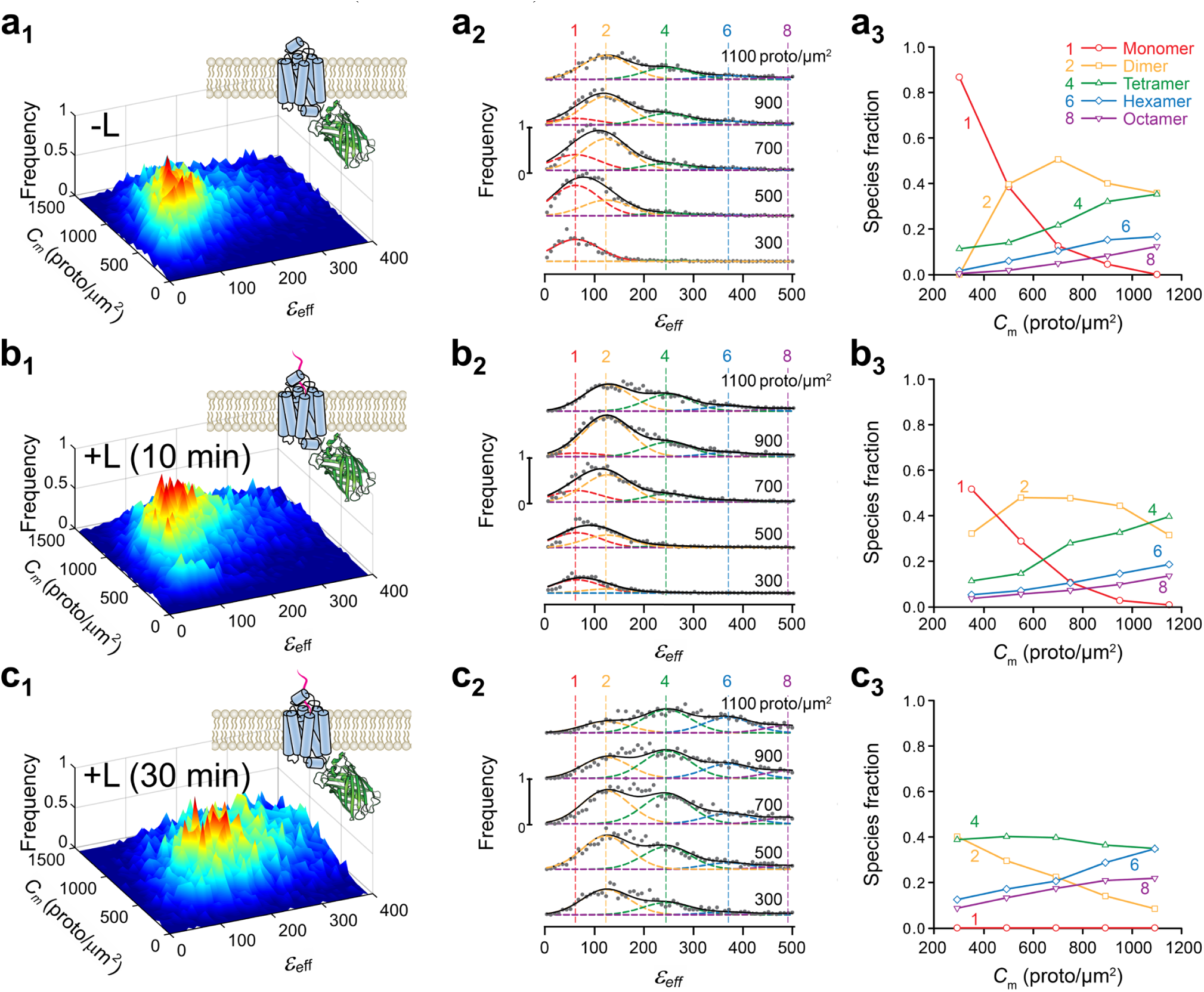
Analysis of effective brightness (*ε*_*eff*_) distributions obtained from two-photon excitation of cells expressing wild-type secretin receptor in the absence of ligand (-L) or after ten-minute treatment with 100 nm ligand (+L) for various treatment times. a1-c1 (column 1), Surface plots of the frequency of occurrence of *ε*_*eff*_ for each concentration of protomers using **(a1)** 13,420, **(b1)** 15,309 and **(c1)** 12,979 total segments to construct the distribution. **a2-c2 (column 2),** Stacks of cross sections through the surface plots in panels a, i.e., frequency of occurrence vs. effective brightness for different concentration ranges; average concentration for each range (in protomer/μm^2^) is indicated above each plot. The vertical dashed lines indicate the peak positions for the brightness spectra of monomers, dimers, etc., obtained from (or predicted by) the simultaneous fitting of the PM-1- and PM-2-mEGFP spectrograms used as standards of brightness (see caption to Figure 2). **a3-c3 (column 3),** Relative concentration of protomers within each oligomeric species vs. total concentration of protomers, as derived from unmixing of the curves in column 2 into different Gaussian components. Samples were as follows: wild-type secretin receptor treated with vehicle (-L) (**row a** of graphs), secretin (+L) for 10 minutes (**row b**), or secretin (+L) for 30 minutes (**row c**).

Such a dramatic effect is perhaps not accidental and may indicate that oligomerization plays a role in signaling. This hypothesis will need to be tested in future studies. At this juncture, however, we know that the observed effect was not induced by cell fixation, since we obtained very similar results for living cells (see Supplementary Figure 4).

We are now ready to clarify why this wealth of information extracted with 2D FIF spectrometry is lost when using averages over large regions of interest, as done in other intensity fluctuation-based methods. The explanation lies in the observation that each ROI contains wide ranges of molecule concentrations and oligomer sizes (see the inhomogeneous distribution of intensities in Figure 1), which has two major consequences, as follows.

i. When a single fluorescence intensity histogram is generated from an entire ROI, image intensity non-uniformities caused by inhomogeneous distributions of oligomer sizes and concentrations broaden the histogram, which results in a larger apparent brightness (see equation 1 in the Methods) than would be obtained for more uniform intensity distributions. This effect is usually so large that it obfuscates any difference between the protein oligomer sizes obtained in, e.g., the case of low vs. high concentrations of molecules, or the absence vs. presence of ligand (see Supplementary Figure 5).
ii. Although the oligomer sizes fluctuate from place to place within the same ROI (due to, e.g., fluctuations in concentration or local physico-chemical properties of the membrane milieu), only an average brightness value is obtained if the histogram is based on all the pixels in an entire ROI. Superficially, this would be justified by the need to reduce the noise in the data by using a large statistical ensemble of pixels, but it has the unintended consequence that it does not reflect correctly the diversity of oligomer sizes present in that ROI. In other words, it is not the noise but rather the oligomer size that is averaged out by using large ROIs.

These two effects clearly justify the need for image segmentation, which captures the local fluctuations in molecular distributions and transfers them to the 2D brightness-concentration spectrograms, which allows for extraction of information on proportions of oligomer sizes and stability in 2D FIF spectrometry.

In conclusion, we demonstrated that 2D FIF spectrometry correctly identifies the oligomer size of various fusion protein constructs and is able to distinguish between differing oligomer sizes formed by a prototypical receptor tyrosine kinase as well as its oligomerization-defective mutant. It also allowed us to uncover a striking ligand-induced shift in the oligomeric size of a class B GPCR, the human secretin receptor, in live as well as fixed cells with exquisite sensitivity and accuracy. Our approach is rather simple and intuitive, and the step-by-step process could be easily learned by experienced and inexperienced researchers alike and be used to produce definitive results within just a few days. As it can be implemented on a variety of imaging platforms and is several orders of magnitude faster than other approaches, FIF spectrometry may be readily used for a variety of fundamental studies in cellular signaling as well as for high-throughput screening of drugs targeting protein-protein interactions.

## Methods

### DNA constructs and cell lines

DNA constructs were made as previously described^38,39^. Monomeric A206K mEGFP construct (PM-1-mEGFP) or a tandem dimer of A206K mEGFP (PM-2-mEGFP) were targeted to the plasma-membrane by adding the palmitoylation-myristoylation sequence, (Met)-Gly-Cys-Ile-Asn-Ser-Lys-Arg-Lys-Asp, at the amino terminus of the A206K mEGFP and the A206K mEGFP tandem.

Stable cell lines expressing receptors of interest were generated as described previously^38,39^. Wild-type and mutant EGFR as well as the plasma membrane targeted monomeric and dimeric constructs were expressed in Flp-In™ T-REx™ 293 cells (Invitrogen); these cells were maintained in DMEM (high glucose) supplemented with 10% (v/v) fetal bovine serum, 100 U ml^−1^ penicillin, 100 mg ml^−1^ streptomycin, 10 mg ml^−1^ blasticidin and 100 mg ml^−1^ zeocin. The wild-type secretin receptor constructs were stably expressed in chinese hamster ovary (CHO) cells; stably transfected CHO cell lines were maintained in Hams F-12 Nutrient Mix (Invitrogen, Paisley, U.K.) supplemented with 5% (v/v) fetal bovine serum, 100 U ml^−1^ penicillin, 100 mg ml^−1^ streptomycin and 500 mg ml^−1^ zeocin. All cells were maintained in a humidified incubator with 95% air and 5% CO_2_ at 37°C.

### Cell preparation for imaging

Flp-In^TM^ T-REx^TM^ 293 cells expressing plasma-membrane targeted monomeric A206K mEGFP constructs (PM1-mEGFP) or dimeric A206K mEGFP constructs (PM2-mEGFP) were plated onto poly-D-lysine-coated 30-mm glass coverslips at a density of 2.5∙10^5^ cells/coverslip. These cells were induced using doxycycline at concentrations between 0.25 and 100 ng∙ml^-1^, in order to achieve various expression levels. After induction, the cells were allowed to grow overnight, and the coverslips were then rinsed and resuspended in HEPES buffer (130mM NaCl, 5mM KCl, 1mM CaCl_2_, 1 mM MgCl_2_, 20mM HEPES, and 10mM D-glucose, pH 7.4), where they were then taken for imaging on a confocal microscope.

Secretin receptor-mEGFP, EGFR-mEGFP, and Tyr251Ala,Arg285Ser EGFR-mEGFP and cells were all grown in Lab-Tek 4-well-chambered cover glasses (Thermo Fisher Scientific, Paisley, U.K.). Samples were either treated with ligand (EGF for EGFR expressing cells and secretin for secretin receptor expressing cells) or vehicle for and then fixed with 4% paraformaldehyde. After fixation, paraformaldehyde solution was removed from the chamber, rinsed multiple times in PBS, and the cells resuspended/imaged in PBS.

### Imaging using single-photon confocal microscopy

Fluorescence images (1024 × 1024 pixels^2^) were acquired using a Zeiss LSM 510 PASCAL EXCITER laser scanning head coupled to a Zeiss Axiovert 200M inverted microscope (Carl Zeiss Microscopy) equipped with a 63× plan apochromat oil immersion lens with a numerical aperture of 1.4. The pixel dwell time was set to 12.8 μs/pixel. Detection of emitted fluorescence was accomplished using a photomultiplier tube (PMT) with settings: gain=850 V, offset=0, and amplifier gain=1. A long pass beam splitter with center wavelength 490 nm along with a long pass emission filter with a wavelength of 505 nm were chosen to efficiently collect the A206K mEGFP emission signal. All samples were excited using the 488-nm line of the 25-milliwatt multiargon laser. The 1/e^2^ laser beam waist radius was estimated by imaging a *z* stack of sub-diffraction-sized 100-nm Tetraspeck fluorescent microspheres (Invitrogen, catalog no. T14792) and found to be *ω*_0_=0.266 μm for radial direction. The contribution of background signal as well as detector shot noise to the measured intensity distributions was determined as described in Supplementary Note 4.

### Imaging using two-photon microscopy

Fluorescence images (800×480 pixels^2^) were acquired using a two-photon optical micro-spectroscope^31,40^ comprised of a Zeiss Axio Observer inverted microscope stand and an OptiMiS detection head from Aurora Spectral Technologies. A mode-locked laser (MaiTai^TM^, Spectra Physics), which generated 100 fs pulses, was used for fluorescence excitation at 960 nm. The excitation beam was focused in the plane of the sample using an infinity-corrected, C-Apochromat, water immersion objective (63×, NA=1.2; Carl Zeiss Microscopy). The optical scanning head (for laser beam scanning) was modified to incorporate a spatial light modulator (SLM) (P1920-1064-HDMI Nematic SLM System, Meadowlark Optics) for adaptive laser beam shaping. A multi-beam array was generated using the SLM and appropriate software, for exciting 40 voxels in the sample simultaneously; the average power for each voxel was 4 mW. The OptiMiS detection head employed a non-descanned detection scheme, in which the emitted fluorescence was projected through a transmission grating onto a cooled electron-multiplying CCD (EMCCD) camera (iXon Ultra 897, Andor Technologies), allowing for the different wavelengths of light emitted by the sample to be separated into wavelength channels simultaneously (i.e. from a single exposure). The spectral bandwidth of the wavelength channels ranged from 450 nm to 600 nm with a spectral resolution of 22 nm. The 1/e^2^ laser beam waist radius was estimated by imaging a *z* stack of sub-diffraction-sized 170-nm PS-Speck Microscope Point Source Kit fluorescent microspheres (Invitrogen, catalog no. P7220) and found to be *ω*_0_=0.316 μm for radial direction. The spatial light modulator (for laser beam shaping), optical scanning head (for laser beam scanning), and EMCCD camera used for image acquisition were controlled by the same computer using in-house custom software written in C++.

### Molecular brightness and receptor concentration determination

The essence of the fluorescence fluctuation spectroscopy methods is that the variance, *σ*^2^, of the distribution of measured intensities is dependent on both the number of photons emitted per second per molecule, i.e. the molecular brightness *ε*, and the average number of particles (or oligomers) within the observation volume, *N*_*oligo*_^41^. The assumption is that both the fluctuations in fluorescence and the detector shot noise follow Poisson statistics. When using analog detectors for signal collection, the molecular brightness, *ε*, can be extracted from the variance of the intensity distribution using the following relation^42^:

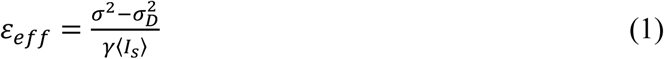

where *ε*_*eff*_ ≡ *Gε* and G is the analog gain in digital levels /photon, 〈*I*_*S*_〉 is the average background corrected measured intensity, *γ* is a shape factor which depends on the shape of the laser PSF as well as the geometry of the sample^43^, and 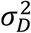 is the variance arising due solely to the detector, which can be obtained from separate measurements on a constant intensity light source (see Supplementary Note 1 for a more detailed derivation).

The total number of protomers within the beam volume, *N*_*proto*_, can be written as a function of the measured intensity, as follows:

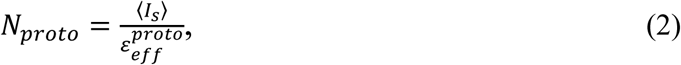

where 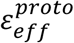 the molecular brightness of a single protomer, which must be determined from applying equation (1) to separate measurements of a calibration sample known to be monomeric. The concentration of protomers, *C*, can then be determined by dividing *N*_*proto*_, by the value for the observation volume:

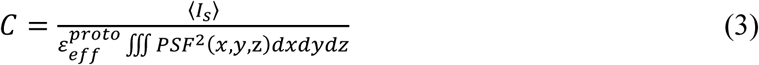

Here PSF represents the laser distribution function of the focused laser beam and ∭ *PSF*^2^(*x*, *y*, *z*)*dxdydz* the volume of the laser beam comprising *N*_*proto*_ molecules (see Supplementary Note 2 for detailed derivation) and was numerically evaluated using a program written in Matlab^TM^ (Matworks Inc.). For measurements of basal membranes of cells expressing membrane proteins, the concentration of molecules in the membrane, *C*_*m*_, becomes *N*_*proto*_ per beam area, which is a particular case of equation (9), given by:

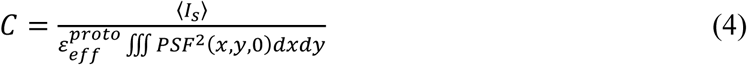

### Description of the data analysis program

A graphical user interface was created that encompasses all the steps needed to quantify oligomeric actions from fluorescence fluctuation data. The software suite is separated into three modules: (1) region of interest (ROI) and segmentation generation, (2) brightness and concentration extraction, and (3) meta-analysis of brightness distributions. Each module is launched by a separate icon in the graphical user interface (GUI) toolbar.

In the first module, 2D fluorescence images are loaded as a stack, and the user selects ROIs using a polygon tool; multiple ROIs can be selected for each loaded image. A segmentation process is implemented which divides each ROI into smaller segments using either a moving square algorithm or a simple linear iterative clustering algorithm (SLIC) (for more details see Supplementary Note 3)^44–46^. The pixel locations for each segment are saved and paired with the corresponding source image. For a comparison of the results obtained with the two segmentation methods, see Supplementary Figure 6. The automatic ROI segmentation not only allowed the conversion into critical information of the inherent average intensity variations from segment to segment, but it also increased the number of data points using only a reasonably small number of manually selected ROIs of 100 or so.

Once all the ROIs are drawn and segments generated, the intensity histogram from each segment is fit with a single Gaussian function to determine the mean and the standard deviation for the intensity distribution of said segment; the algorithm executed in the fitting process incorporates the Nelder-Mead method ^47,48^. Using the mean and standard deviation obtained from the Gaussian fitting along with the corrections for the shot and background noise (see Supplementary Note 1), the effective brightness, *ε*_*eff*_, and concentration of the corresponding segment is found by applying equations (2) and (4), respectively. The entire procedure of fitting starting at intensity histogram calculation and ending in the calculation of segments average effective brightness and concentration of the studied molecule is performed after a click of a single button. Once *ε*_*eff*_ and concentrations have been found for each segment, multiple tools for visualization of the brightness distributions as a function of concentration are included. The first visualization tool creates a 3D surface plot of the concentration vs. *ε*_*eff*_, as seen in Figure 2.a1-b1. The second visualization tool allows partitioning each 3D plot into one dimensional brightness spectrograms for several chosen concentration ranges and plots the histograms one on top of one another in order of increasing concentration, as is seen for instance in Figure 2.a2-b2.

The *ε*_*eff*_ distributions for various concentration ranges are further analyzed in the third module where the distributions are fit with a sum of multiple Gaussian functions 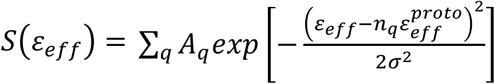; this fitting, again, was performed using the Nelder-Mead method. The means of each Gaussian used in the fitting, 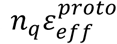, are all linearly related and set equal to a multiple of the monomeric molecular brightness, 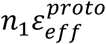. The center of each Gaussian then corresponds to the expected peak of either monomers, dimers, or and various higher order oligomers, depending of the multiplication factor used, i.e. *n*_1_ = 1 for monomers, *n*_2_ = 2 for dimers, *n*_4_ = 4 for tetramers, *n*_6_ = 6 for hexamers, *n*_8_ = 8 for octamers, and *n*_10_ = 10 for decamers. The monomeric molecular brightness, 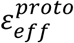, can be found by applying the same software tools to images acquired from a standard monomeric sample and making a single bin of concentration values. For a given concentration range, the relative amplitudes of the *i*^*th*^ Gaussian used in the fitting, *n*_*i*_*A*_*i*_/ ∑_*q*_*n*_*q*_ *A*_*q*_, indicates the fraction of total protomers that the corresponding oligomeric species comprises. Plots of the species fraction values for each oligomer size, obtained from the relative amplitudes of the Gaussian fittings, are shown in Figure 2a3-b3.

### Computer program availability

Software used for data analysis described in this work is available upon request.

## Data availability

Fluorescence images and ROI files used to generate the FIF spectrograms in this study are available upon reasonable request.

## Acknowledgments

Mrs. Annie Eis for assistance with protein construct design and purification, and Dr. Larry Miller, Mayo Clinic, Arizona, for provision of the Secretin receptor–mEGFP expressing cells that we used to develop the current cell lines. We also thank Marwan McBride and Alexander Klug for assistance with data analysis. This work was partly funded by the National Science Foundation grant PHY-1626450 (awarded to V.R.), the UWM Research Growth Initiative grants 101X333 (to V.R.) and 101X340 (to I.V.P.), and Medical Research Council U.K., grant number MR/L023806/1 (to GM).

## Author contributions

M.R.S. prepared samples, performed two-photon microscopy measurements, designed and implemented algorithms, performed data analysis, and participated in manuscript writing. G.B. implemented data fitting algorithms, wrote the computer program for data reduction and analysis. R.J.W. and J.D.P. designed the DNA constructs and cells lines, prepared the samples, and performed confocal microscopy measurements. D.B. participated in sample preparation and two-photon microscopy measurements. I.V.P. designed DNA constructs and supervised work on expression and purification of fusion fluorescent proteins. G.M. designed DNA constructs, supervised the work on development of cells lines and confocal microscopy imaging, and participated in manuscript writing. V.R. conceived and designed the study, generated algorithms, participated in data analysis, wrote the manuscript together with M.R.S. and G.M. and with input from G.B. and I.V.P., and supervised the project.

## Additional Information

**Supplementary information** consisting of figures and notes is attached.

**Correspondence and request for materials** should be addressed to V.R.

